# Spurious off-target signals from potential lncRNAs by 10X Visium probes

**DOI:** 10.1101/2022.09.25.509336

**Authors:** P. Prakrithi, Juwayria, Deepali Jain, Prabhat Singh Malik, Ishaan Gupta

## Abstract

Spatial transcriptomics has revolutionized molecular profiling of tissues in a spatial context, especially in the study of cancer heterogeneity. 10X Genomics facilitates spatial gene expression profiling platforms to help work with fresh-frozen (FF) and formalin fixed paraffin embedded (FFPE) tissues. FF analysis is based on polyA capture of RNAs while FFPE analysis uses a pre-designed set of probes to capture transcripts of coding genes. Previously, we used FFPE spatial data as a ‘negative control’ in a study to identify novel non-coding RNAs in FF data. Interestingly, we find and report that certain target probes used in FFPE show off-target signals from lncRNAs. The Space Ranger pipeline of 10X Visium counts the expression of these potential off-targets to be that of the corresponding target gene, some of which have known implications in cancer and its diagnosis. Therefore, relying on this technology is not ideal to investigate expression of the genes reported in this study. We hereby recommend excluding those genes in any downstream analysis of FFPE datasets and to design probes with better specificity, considering the sequence similarity between genes and non-coding RNAs.

## Detection of spurious off-target signals from potential lncRNAs by 10X Visium probes

Over the past few years, the world has seen a lot of technological advancements in the field of transcriptomics. Understanding transcriptomics throws light on the dynamic molecular state of cells and helps uncover the genetic basis of various diseases like cancer. Since tumours are heterogeneous, and involve various cell types and their interaction with each other, it is important to profile the transcriptome without losing the spatial context. The expression profiles from bulk RNA-sequencing would not give us such information. This has been made possible with the recently developed spatial transcriptomics sequencing (ST-Seq). ST-Seq provides the spatial information, i.e where the cells and gene expression signals come from in a tissue section and helps understand cellular interactions [1]. ST-Seq uses histological images where the tissue is sliced and fixated to a microscopic slide with oligos incorporated on the slide’s surface. These capture RNA from the tissue which is further sequenced to understand the transcriptomic signature.

10X Genomics has been pioneering in spatial gene expression profiling through the development of novel spatial platforms marketed under the name ‘Visium’. 10X Visium offers two protocols (i) for fresh frozen (FF) tissue samples based on PolyA-tail capture of RNA (ii) for Formalin-fixed paraffin-embedded (FFPE) samples . However, RNA from FFPE tissue blocks is generally degraded resulting in the disruption of PolyA tails. 10X Visium thus offers a probe-hybridization based protocol, to overcome this limitation. Each probe has an LHS and RHS fragment (termed as probe pairs) to efficiently capture coding RNA molecules. Once bound to the target RNA, the probe pairs are ligated with each other.

For FFPE samples, the 10X Visium human transcriptome probe set v1.0 provides a panel of 19,144 gene_ids targeted by 19,902 probes. 1,201 (6.3%) of these gene_ids (targeted by 1,272 probes) are excluded by default due to predicted off-target activity to homologous genes or sequences. These probes are marked with ‘FALSE’ in the ‘included’ column of the probe set reference CSV. A gene with at least one probe with predicted off-target activity will be filtered out. 17,943 gene_ids (targeted by 18,630 probes) are present in the final filtered output of the Space ranger pipeline that generates a count matrix for genes per spot.

The ligation events of the pair of probes targeting each gene are counted using Space ranger’s probe aligning algorithm. The reads are aligned to the probe set reference and assigned to the genes they target and the aligned bam file is tagged with the probe and gene ID.

**Figure 1:**
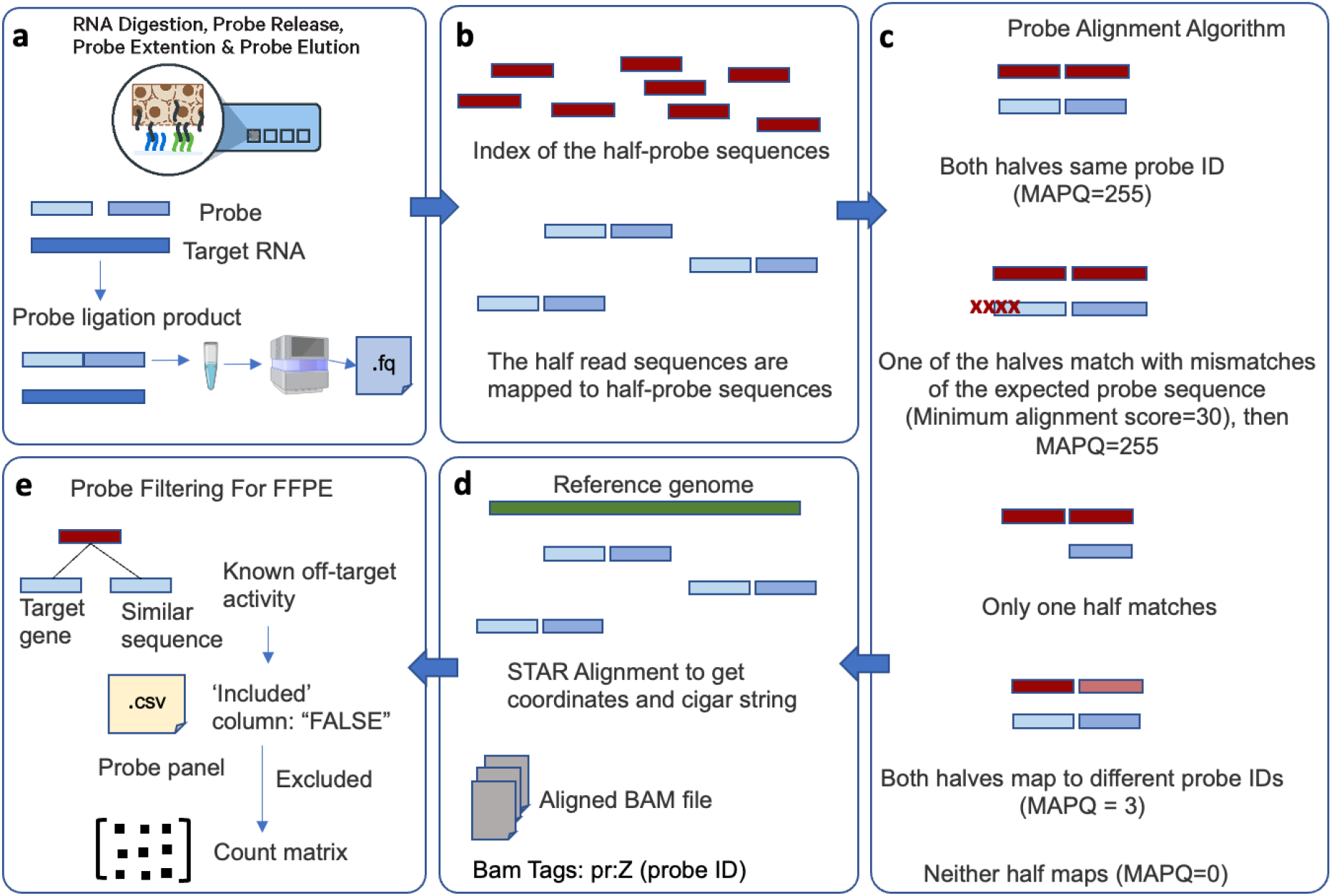
Overview of the 10X Visium Space ranger pipeline (a) The RNA-hybridized probe ligation products are captured on the Visium slide after the tissue permeabilization and are sequenced to produce raw FASTQ files (b) Space ranger pipeline uses a probe alignment algorithm where each half-read is mapped to every half-probe sequence and (c) assigns quality scores to the transcript reads based on their alignment to the probe sequences (d) Alignment to the reference genome with STAR is done to identify the coordinates and cigar strings of the probe-aligned reads which are tagged with the probe and corresponding gene IDs in the output BAM file (e) Probes with known off-target activity are labeled ‘FALSE’ in the probe sheet and are excluded by default while generating the gene count matrix

10X Visium FFPE spatial transcriptomics data of five public datasets and two in-house were used as a control in a study to identify potential novel non-coding transcripts in FF tissue samples using a pipeline that identifies transcriptionally active regions (TARs) and classifies them as annotated (aTARs) and unannotated (uTARs) based on their overlap with existing gene annotations[2].

**Table 1:**
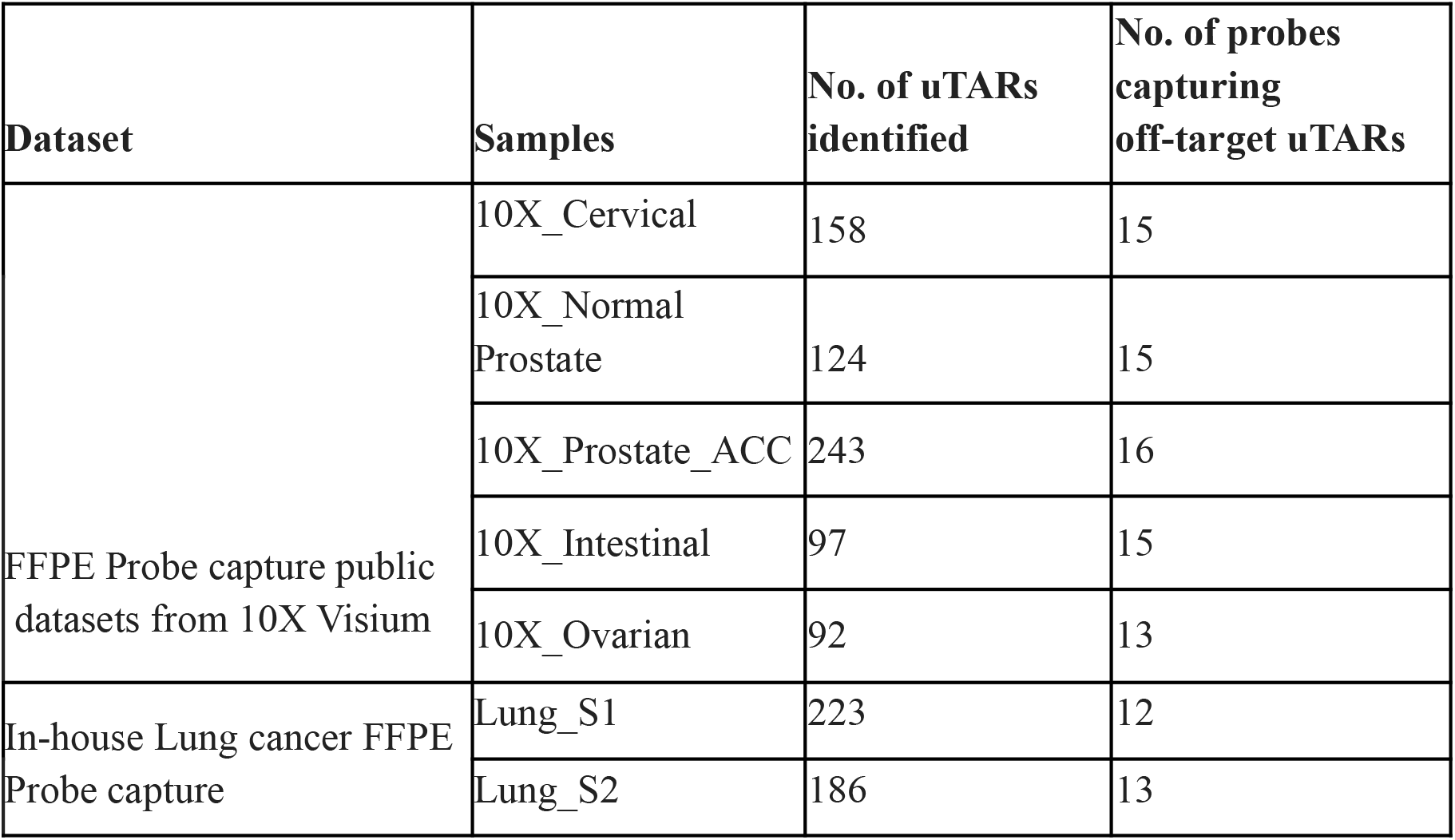
Number of potential non-coding RNAs identified.

| Dataset | Samples | No. of uTARs identified | No. of probes capturing off-target uTARs |
| --- | --- | --- | --- |
| FFPE Probe capture public datasets from 10X Visium | 10X_Cervical | 158 | 15 |
|  | 10X_Normal Prostate | 124 | 15 |
|  | 10X_Prostate_ACC | 243 | 16 |
|  | 10X_Intestinal | 97 | 15 |
|  | 10X_Ovarian | 92 | 13 |
| In-house Lung cancer FFPE Probe capture | Lung_S1 | 223 | 12 |
|  | Lung_S2 | 186 | 13 |

The 10X Visium FFPE protocol sequences a probe that is complementary to the mRNA it is hybridized with. Due to this we found a higher number of TARs antisense to the gene annotations. This conflict in the orientation should be considered before running the uTAR-Seq pipeline since the antisense TARs would be considered as uTARs by default when directionality is considered, rather than them being considered aTARs.

Although one would not expect to capture novel transcripts from this FFPE protocol since the capture is targeted, hundreds of uTARs were still identified in gene deserts. About 42% (102) of these uTARs were found to overlap with lncRNAs from public databases such as FANTOM[3] and LncExpDB [4] (Fig. 2a), confirming that these captured uTARs are not false positives and are predicted lncRNAs. To identify how these off-targets were captured, the probe sequences and the read sequences were analyzed. Hamming distances were calculated using the R package DescTools [5]. 22 probe sequences matched exactly with the uTAR sequences while majority of the uTARs identified have a Hamming distance of 1 and 2, indicating high sequence similarity and hence captured by the probes meant to target coding genes. These were 117 probes capturing upto 243 uTARs. It was also seen that in the bam files generated by the 10X spaceranger pipeline, these off-target reads were tagged with the genes intended to be captured by the probes. The positions of such reads and the position of the gene with which the reads were tagged was contradictory.

**Figure 2:**
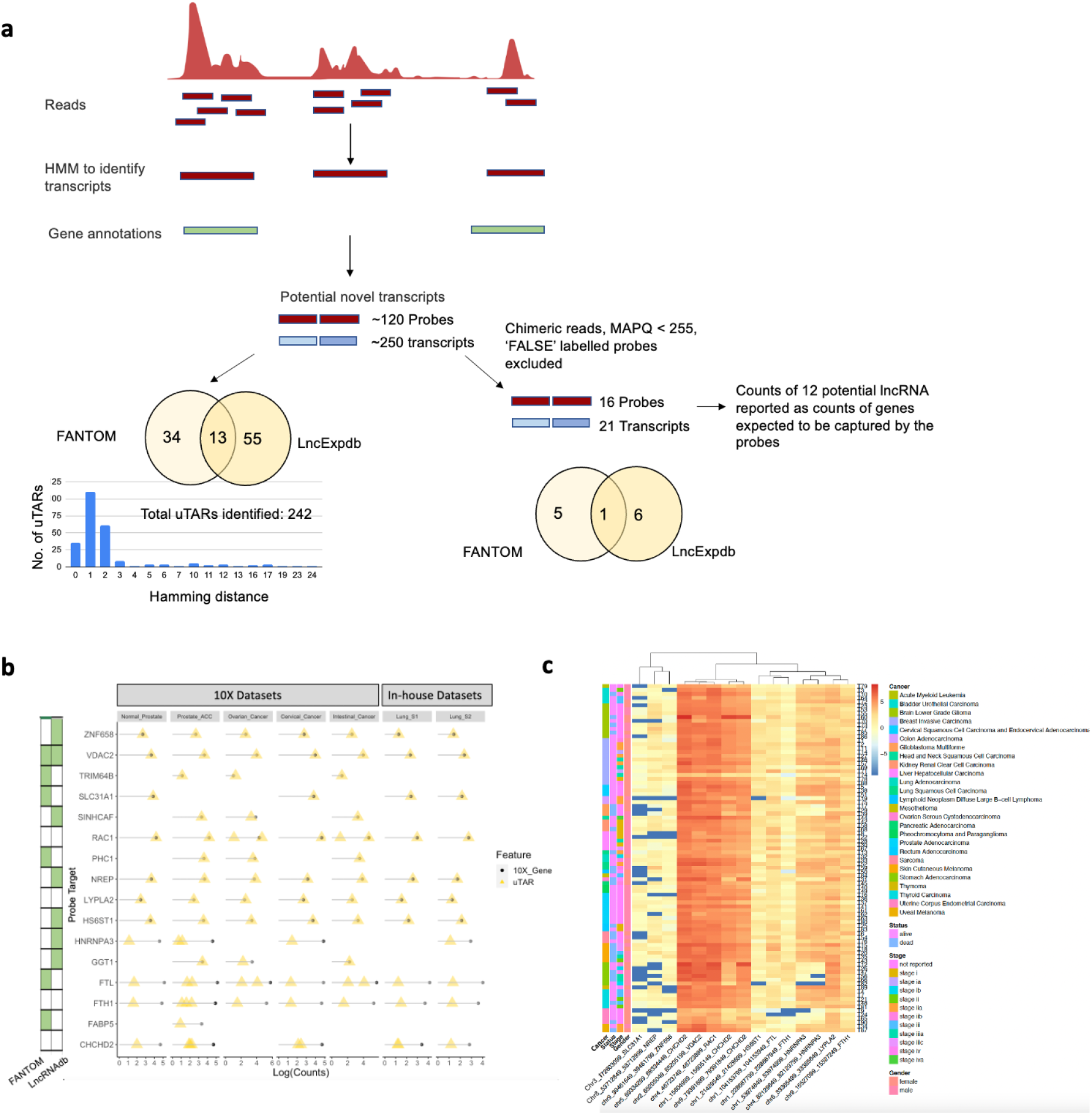
(a) The transcriptionally active regions identified using a HMM identified transcripts outside gene boundaries and their overlap with reported lncRNAs before and after applying filters (b) Probes picking up off-targets, the counts of which are represented as gene counts by 10X Space Ranger as shown by the same counts of the gene and off-target uTAR, some of which are reported lncRNAs in public databases, indicating that these genes should be excluded in preceding analyses (c) Heatmap showing the coverage (log) of the off-target regions from TCGA samples

To be more confident about the spurious signals, the chimeric transcripts, the reads not confidently mapped to probe IDs (MAPQ<255), those tagged with multiple probe IDs and the probes labeled ‘FALSE’ in the ‘included’ column of the probe set .csv file were filtered out. This still resulted in 21 probe-transcript pairs, the probes of which correspond to 16 genes. Of these 21 off-target transcripts, 13 overlapped with lncRNAs from public databases (Table 2). Moreover, it was seen that the counts generated by the 10X Space ranger pipeline for 12 genes that were expected to be captured by the probes were exactly the counts of the 12 potential off-target transcripts, while for the rest, the off-target counts had a minor contribution to the 10X gene counts. The results were replicated in all the samples analyzed, even in the low coverage in-house lung cancer datasets (Fig.2b). BLAT searches of the sequences also confirmed their off-target positions. Furthermore, expression of these regions were also observed from the coverage data (BigWig files) from RNA-Seq datasets of The Cancer Genome Atlas (TCGA) (Fig.2c).

**Table 2:**
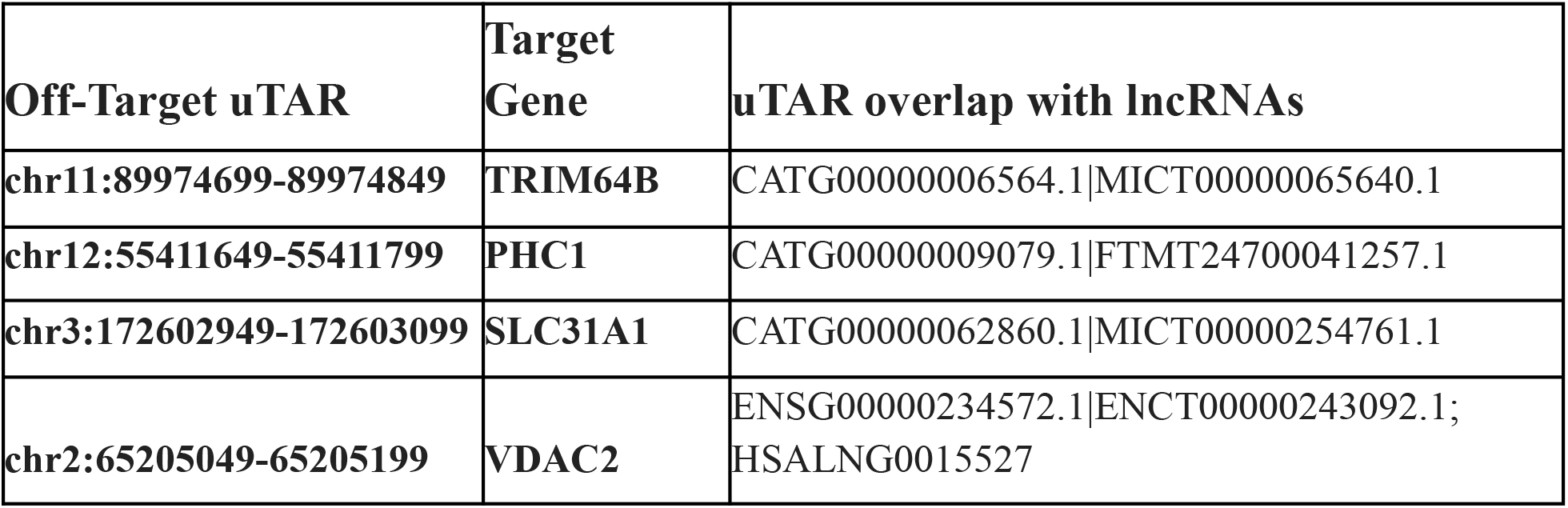
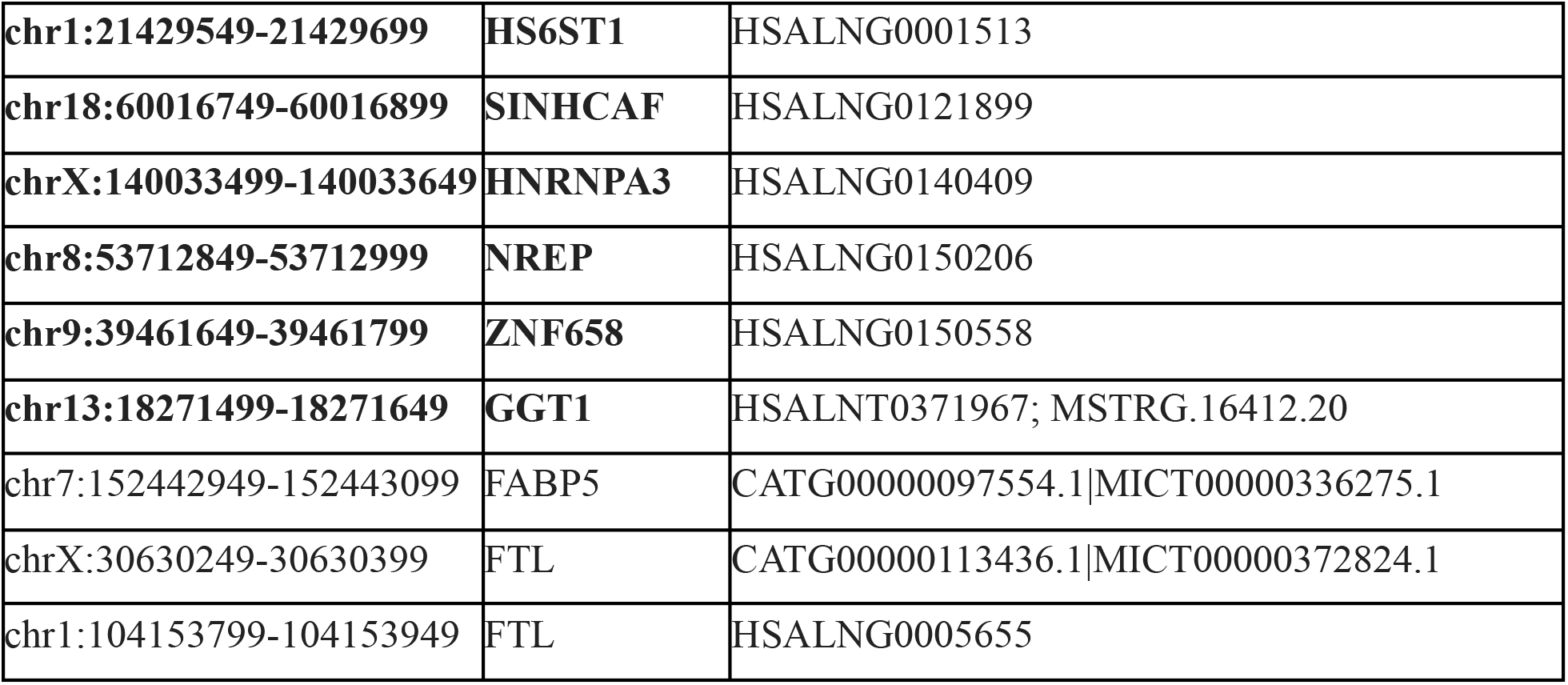
Off-target uTAR captured by probes designed for corresponding target genes and the lncRNA IDs from public databases that overlap with the uTARs.

To see how these spurious off-target signals affected the gene expression counts reported by 10X spaceranger, we performed a correlation analysis between the counts reported by the uTAR-Seq pipeline and 10X. This was done for (i) aTAR counts corresponding to a gene Vs gene counts reported by Space ranger and the (ii) aTARs along with the off target counts tagged with the gene Vs the gene counts from Space ranger. The correlation improved in the latter case, suggesting that 10X might be wrongly reporting the off-target signals as gene counts. This suggests that counts from such probes must be excluded in the downstream analyses due to their off-target binding potential.

The genes for which Space Ranger reports the counts potentially picked up from lncRNAs have been implicated in cancer. For example, the gene *GGT1* has been used to study the complexity of nephrons in kidney cancer using the Visium platform. However, a uTAR (chr13:18271499-18271649) that is a lncRNA reported in LncRNABook[4] is non-specifically captured by the probe designed to capture this gene. The expression of this uTAR has been reported by the 10X pipeline to be that of the *GGT1* gene and hence cannot be relied upon. Similarly, genes like *SINHCAF* that has been implicated in the regulation of hypoxia in esophageal cancer[6]. Moreover, the outer mitochondrial membrane voltage-dependent anion channel (VDAC) - MCL-1 interaction has been shown to increase mitochondrial Ca^2+^ uptake and reactive oxygen species generation in lung cancer cells and hence a marker for hypoxia[7]. Similarly, NREP overexpression promotes migration and invasion, in the activation of cancer associated fibroblasts/ EMT and is used as a prognostic biomarker [8]. RAC1 expression could be used to assess tumorigenic progress in colorectal cancer [9]. The gene HNRNPA3 is used to monitor lymph node metastasis in bladder cancer [10]. The expression of such genes (Table 2, Figure 2b) reported by 10X Visium that might be coming from the uTARs sharing high similarity with these probe sequences could mislead clinical interpretations. Thus the genes reported in this study must be excluded while analyzing FFPE spatial transcriptomics datasets. Since during the probe designing, the off-target binding potential was only analyzed among genes and not other sequences like lncRNAs, these probes must be redesigned considering all the other known non-coding transcripts apart from the coding transcripts for more specific capture of the targeted genes.

## Methods

### Datasets

Public data of cervical cancer, normal and cancerous prostate tissues, intestinal and ovarian cancers from the 10X Genomics site were used for the analysis (https://www.10xgenomics.com/resources/datasets/) [11–15]. Moreover, FFPE tissue biopsies of in-house lung cancer samples from the All India Institute of Medical Sciences (AIIMS), which were of low coverage were also analyzed.

### Pipeline

A recently published pipeline by Wang et. al, 2021 [2] has been used to identify transcriptionally active regions from 10X Visium spatial datasets. The FFPE datasets were used as a control as one wouldn’t expect to find potential novel non-coding RNAs, since the capture is targeted by probes complementary to coding genes. The pipeline uses a Hidden Markov Model to scan the aligned transcriptome for transcriptionally active regions, which are then overlapped with existing gene annotations (Gencode v36, GRCh38/hg38). The TARs that do not overlap the annotations, irrespective of the strandedness, are labeled as unannotated transcriptionally active regions (uTARs). These could be potential novel non-coding RNAs. They are further confirmed by overlapping them with annotations of long non-coding RNAs from dedicated consortia like FANTOM and LncExpdb.

## Data and Code availability

All codes used in the analysis can be found at https://github.com/Prakrithi-P/FFPE_probes_off_target/

## Author contributions

I.G. conceptualized the study. P.P. performed all analyses, generated the figures and wrote the manuscript. All authors revised the manuscript and approved it for publication. J contributed to the sequencing. P.S.M. and D.J. were involved in ethical clearances, patient sampling and clinical annotations of the in-house datasets used.

